# Regional Differences in Progenitor Consumption Dynamics Shape Brain Growth during Development

**DOI:** 10.1101/2023.08.21.553891

**Authors:** Natalia Baumann, Robin Wagener, Awais Javed, Philipp Abe, Andrea Lopes, Adrien Lavalley, Daniel Fuciec, Elia Magrinelli, Sabine Fièvre, Denis Jabaudon

## Abstract

Developing mammalian brains are characterized by disproportionate growth of the forebrain compared to other regions. How this localized expansion occurs is, however, largely unknown. To address this, we identified region-specific neurogenic patterns by creating a single-cell-resolution birthdate atlas of the mouse brain (https://www.neurobirth.org). We report that in forebrain regions, neurogenesis is sustained compared to the hindbrain, where neurogenesis is transient and limited to early brain development. Sustained forebrain neurogenesis reflects lengthened cell cycle and reduced consumptive divisions of ventricular zone progenitors, resulting in a preserved germinal cell pool. Using single-cell RNA sequencing, we identify functional molecular programs of ventricular zone progenitors that spatially and temporally regulate progenitor cycling properties, including through loss-of-function of the forebrain-enriched mitochondrial membrane protein Fam210b. These results reveal a parsimonious mechanism to locally regulate neuronal production, in which the time window during which progenitors generate cells is a critical determinant of region-specific brain expansion.

## INTRODUCTION

Vertebrate brains share a common anatomical blueprint in which three main regions, the hindbrain, midbrain, and forebrain develop during embryogenesis to form the circuits underlying specific functions. While the hindbrain and midbrain (together, the brainstem) control and regulate vital body functions such as heart rate, breathing, sleeping, and eating, the forebrain, and particularly the neocortex in mammals, subserves higher-order functions required for multisensory processing and executive planning. Although these three brain subdivisions are present in all vertebrates, their relative importance varies from one species to another^1^. Birds and mammals have strikingly large forebrains and, in mammals, expansion of the neocortex — the most rostral part of the forebrain, which is particularly developed in primates — underpins the remarkable cognitive abilities of these species.

In all vertebrates, the brain and spinal cord originate from a single layer of neuroepithelial cells that form an initially smooth neural tube. During embryogenesis, this tube thickens locally, causing its curvature and caudo-rostral parcellation into three primary vesicles that will form the hind-mid- and forebrain. The relative proportions of these three segments determine the final shape and size of the brain. In mammals, consistent with the large size of the forebrain compared to the brainstem, the rostral part of the neural tube expands dramatically^2,3^. The neurons that populate each of these three segments are born either directly from progenitors located in the ventricular zone (VZ) lining the ventricles — these cells are called apical progenitors (APs) — or from so-called intermediate progenitors (IPs), which are born from APs but are located in the subventricular zone (SVZ), i.e. away from the ventricles, and that are thought to amplify neuronal production from APs^4^. While the molecular mechanisms underlying the parcellation of the neural tube into three primordial segments are relatively well understood^5^, how their respective expansion is regulated during embryogenesis remains largely unknown and is the topic of the current study.

At the level of progenitors, differences in brain segment size may reflect differences in the onset of the time of cell divisions, in the rate of cell divisions, in cell survival, or a combination thereof. While the timing of the birth of various brain structures has been exquisitely characterized by the seminal work of Joseph Altman and Sherley Bayer (see neurondevelopment.org for a compilation of their publications), a systematic, cellular resolution, unbiased assessment of the coordinated date and pace of birth of cells across the whole mouse brain is still lacking. Here we provide such data, which we show can be used as a resource to identify the mechanisms underlying the differential expansion of brain regions during development and the related assembly of these regions into circuits.

In this study, we used mouse embryogenesis as a model to investigate the differential expansion of brain regions during mammalian development. First, we generated a single-cell resolution atlas of the developing brain to establish the time and duration of birth of all structures of the mouse brain. We find that, in contrast to mid- and hindbrain structures, whose generation is transient and limited to early stages of brain development, the generation of forebrain structures is sustained and extends into late embryogenesis, suggesting differences in the neurogenic potential of progenitors in these regions. Investigating this possibility, we reveal that sustained forebrain neurogenesis reflects lengthened cell cycle and reduced consumptive divisions of ventricular zone progenitors in these regions compared to the hindbrain, resulting in a longer time window during which cells can be born. In the second phase of this study, to investigate the molecular mechanism underlying these differences in AP behavior, we generated a single-cell transcriptomic atlas of VZ progenitors along the developing neural tube, allowing us to identify independent transcriptional programs controlling the temporal and spatial diversity of AP identity. Finally, we demonstrate the functional relevance of such programs in regulating region-specific cell-cycle properties, by manipulating the expression of the mitochondrial membrane protein FAM210B, which is specifically expressed in slow-cycling late forebrain APs: repression of this gene reprogrammed the cycling behavior of forebrain APs towards that of their hindbrain counterparts. Together, these findings shed light on the cellular mechanisms underlying the differential expansion of brain regions during development and related circuit assembly.

## RESULTS

### Generating a single-cell resolution atlas of time of birth across the entire mouse brain

To identify neurons born at single time points across the entire developing embryonic brain, we generated two datasets using two complementary fate-mapping strategies (**Figure 1A**). In Dataset 1, we used ethynyldeoxyuridine (EdU) injections to birthdate-label progenitors across the brain—EdU is injected intraperitoneally in the pregnant dam and, as a thymidine analog, is incorporated into the DNA of S-phase cells^6^— and in Dataset 2, we used FlashTag (FT) fate-mapping to birthdate-label dividing APs—FT is injected into the cerebral ventricles of the embryos and specifically labels mitotic APs since they are located juxtaventricularly^7^. Hence, while FT birthdate-labels AP-born cells, EdU birthdate-labels AP- and IP-born cells^7,8^. We performed injections on sequential embryonic days (E) between E10.5 and E17.5, essentially covering the period of brain neurogenesis. In Dataset 2, to ensure that labeled cells underwent a single round of division (i.e., were directly born from APs), we combined the FT pulse with the chronic delivery of bromodeoxyuridine (BrdU) via an intraperitoneal osmotic pump inserted in the pregnant dam (**Figure S1A**). This allowed us to unambiguously identify cells born directly from APs as FT^+^BrdU^−^ cells (i.e. cells that were born at the time of FT injection but that never underwent a subsequent cell division in the presence of BrdU) (**Figure S1B**)^9^.

**Figure 1.**
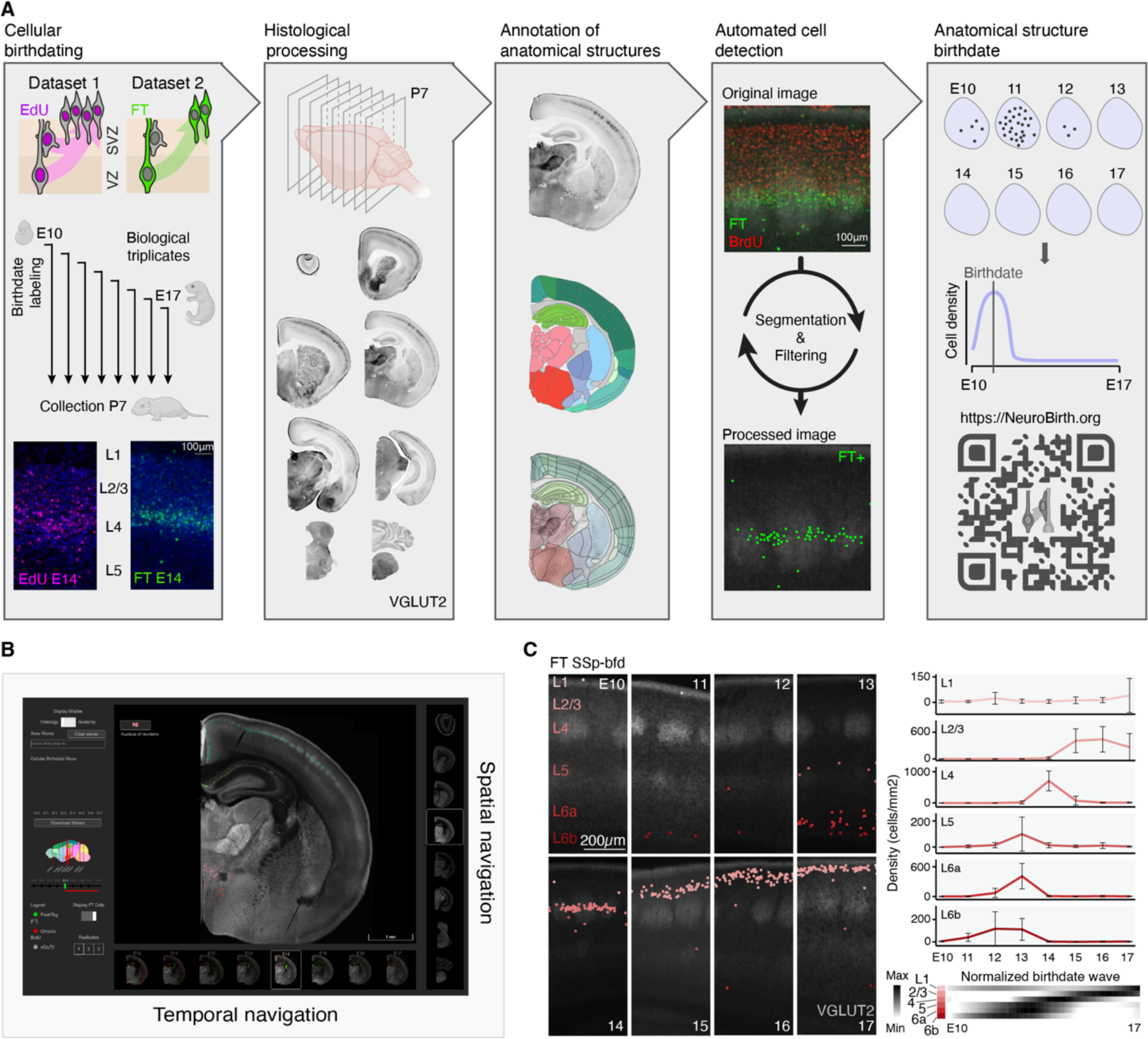
Generating a single-cell resolution atlas of time of birth across the entire mouse brain. (A) Schematic illustration of the experimental and analytic pipeline. EdU (Dataset 1) or FT (Dataset 2) pulse-injections were performed from E10.5 to E17.5, brains were collected at P7, sectioned and stained. Anatomical structures were manually annotated using the Allen Brain Atlas, images were automatically analyzed and EdU^+^ or FT^+^ cells were counted by age for each anatomical structure to build structure-specific birthdate-waves. (B) Screenshot from the https://www.neurobirth.org webtool. (C) Detected FT^+^ cells with VGLUT2 counterstaining in the barrel field somatosensory cortex (SSp-bfd, left) and cell densities of corresponding cortical layers, error bars show the standard deviation across replicates (right).

We collected brains on postnatal day (P)7, i.e., once cellular migration is largely complete, coronally sectioned them, and stained these sections using immunofluorescence for EdU (Dataset 1), or FT and BrdU (Dataset 2). To delineate brain structures according to anatomical landmarks, we used DAPI nuclear counterstaining and immunorevealed for the vesicular glutamate transporter 2 (VGLUT2). For each injection age, eight different rostro-caudal coronal levels were collected from three different animals, resulting in a dataset of 384 images from a total of 128 animals (62 for Dataset 1 and 66 for Dataset 2). The sections were then manually delineated and annotated into 385 regions using the nomenclature and hierarchy of the mouse Allen Brain Atlas for each of the triplicates (www.brain-map.org and see Methods). Of note, we did not analyze data for cerebellar structures, since a substantial fraction of their development occurs postnatally^10^, i.e. beyond the time points included in this study. Using automated cell detection and semi-supervised cell filtering, EdU^+^ cells (Dataset 1) or FT^+^BrdU^−^ cells (hereafter abbreviated FT^+^ for simplicity; Dataset 2) were counted and registered according to injection age and anatomical location, allowing us to quantify structure-specific cell birth dynamics across the entire mouse brain (**Figure 1A**, **Figure S1A-C**, **Methods**). All photomicrographs and associated data were integrated into an online interactive website freely accessible to the community (https://www.neurobirth.org) (**Figure 1B**).

To compare time and pace of neurogenesis across all 385 annotated brain anatomical structures, we developed a common framework in which we computed birthdate “waves”. We segmented embryonic development between E10.5 and E17.5 into 71 epochs (from E10.5 to E17.5 by 0.1 increments) to create a semi-continuous axis (a “pseudotime”) along which we plotted average labeled cell densities across replicates (**Figure 1A**, right). The “birthdate” of a structure was defined as the weighted average of the wave distribution. We validated this experimental and analytic pipeline by assessing the date of birth of well-characterized brain structures. As an example, in the barrel field of the primary somatosensory cortex (SSp-bfd), Datasets 1 and 2 both revealed a classical neuronal inside-out pattern of laminar birth, in which cells positioned in deep layers were born early (from E11.5 to E13.5) while cells of the superficial layers were born later (from E14.5 to E17.5), (**Figure 1C**, **Figure S1E**). Cells detected overwhelmingly represented neurons at all ages and across brain regions, as indicated by cell-type and region-specific analysis of identity from an available developmental dataset^11^ (**Figure S1D**). Taking advantage of this unique dataset, we thus sought for region-specific features of neurogenesis in the developing brain.

### Comparison of EdU and FT birthdate-labeling reveals brain-wide spatio-temporal patterns of direct vs. indirect neurogenesis

One possible mechanism for making larger brain structures is the generation of IPs that amplify neuronal output from APs through indirect neurogenesis^4,12^. While IPs have been well described in the neocortex, the extent to which they are present across brain structures is less clear. To examine patterns of direct vs. indirect neurogenesis across the whole mouse brain, we compared results obtained with EdU labeling (Dataset 1) with those obtained with FT labeling (Dataset 2). EdU labels both APs and IPs while FT only labels APs^7^; consequently, regions with fewer FT^+^ cells than EdU^+^ cells have prominent indirect neurogenesis, while regions with comparable numbers of FT^+^ and EdU^+^ cells have prominent direct neurogenesis. Using this principle, we identified two major regions which, in addition to the neocortex, displayed striking differences in birthdate between Dataset 1 and Dataset 2: the striatum and the thalamus (**Figure S1F-M**), suggesting that a significant number of cells are produced outside of the VZ (i.e in the subventricular zone) in these regions. In the striatum, the difference emerged at E15.5 (**Figure S1F-H**) while in some nuclei of the thalamus (the ventral posteromedial nucleus (VPM) and the posterior nucleus (PO)), FT^+^ cells were not detected after injection at any embryonic age although EdU^+^ cells were present, suggesting that these nuclei are generated mostly through indirect neurogenesis (**Figure S1J-L**). Confirming abventricular cell divisions, and in line with previous work, immunofluorescence revealed cells expressing the mitotic cell markers KI67 or PH3 outside of the VZ at E15.5 in the ganglionic eminences^13,14^, (**Figure S1I**) and at E12.5 in the developing thalamus^15^ (**Figure S1M**). Hence, our datasets highlight differential patterns of direct and indirect neurogenesis across diverse structures of the mouse brain.

### Brain-wide analysis of cellular birthdates reveals region-specific developmental dynamics in the forebrain and hindbrain

Leveraging the birthdate-waves described earlier, we calculated the birthdates of all 385 annotated anatomical structures to explore neurogenesis timing and pace across the whole brain (**Figure 2A,B**). This analysis revealed that brainstem structures primarily develop during early embryonic stages, with 80% forming before E14 (Hindbrain (HB): E13.0; Midbrain (MB): E13.3; Interbrain (IB): E13.9) (**Figure 2B**). In contrast, forebrain structures are generated over a more extended time window, beginning as early as in the brainstem but continuing throughout the time course of our study: it is not until E16 that 80% of forebrain structures are formed (Cortex (CTX): E14.9; Cerebral nulei (CNU): E16.2; **Figure 2B**, right). FT birthdating in Dataset 2 confirmed this prolonged formation phase for forebrain structures, indicating that prolonged generation of forebrain structure applies both to directly- and indirectly-born cells (**Figure S2A-D**).

**Figure 2.**
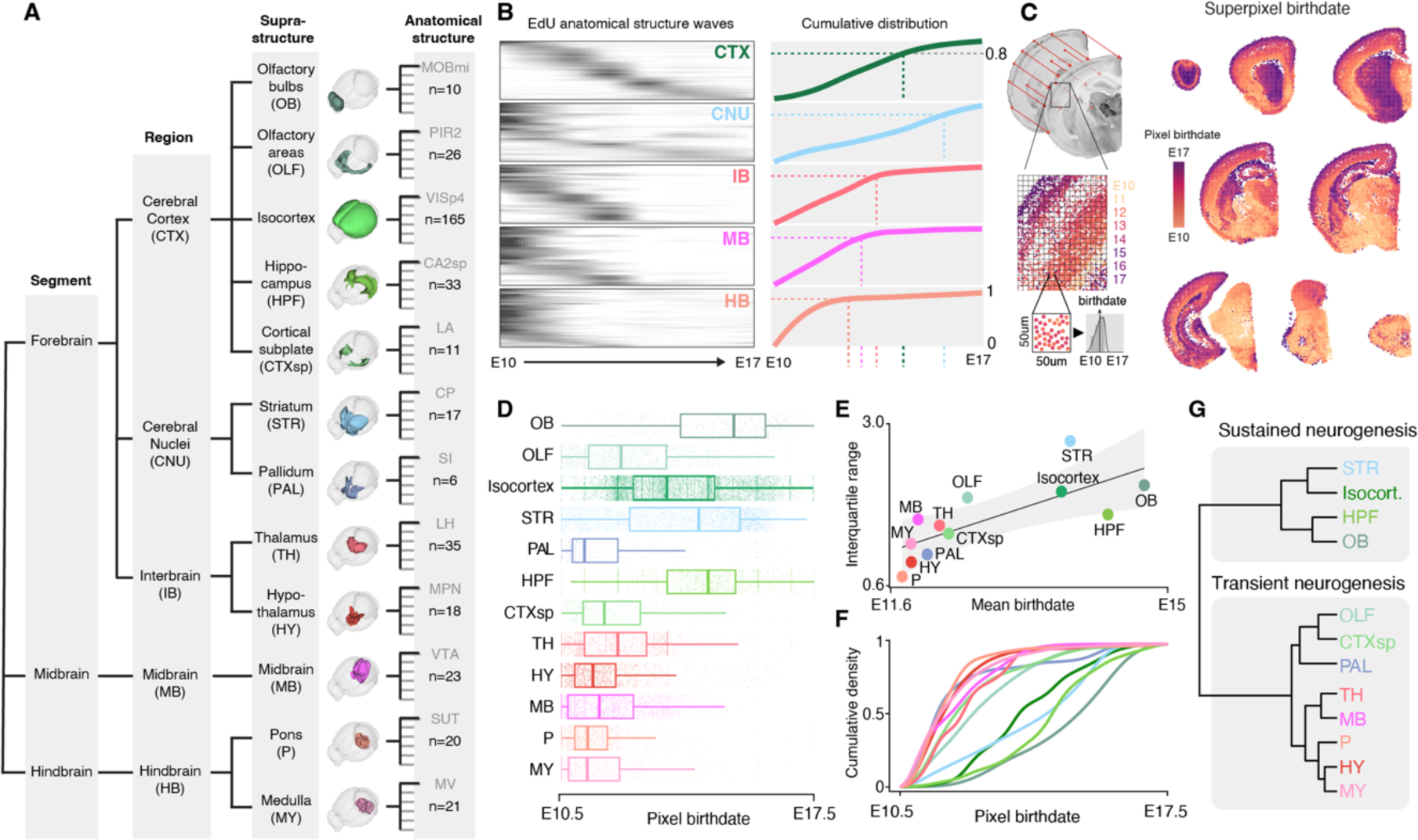
Brain-wide analysis of cellular birthdates reveals rostral sustained-neurogenic and caudal transient-neurogenic regions. (A) Hierarchical tree of mouse brain anatomy highlighting segments, regions, supra-structures subdivision and examples and number of anatomical structures, using the Allen Brain Atlas nomenclature. (B) EdU birthdate-waves for all anatomical structures grouped by regions (left) and cumulative distribution of birthdate-waves (right). The time to reach 80% of born cells is highlighted by the dotted lines. (C) Image alignment allows projection of birth-dated cells onto a common space. The common section is divided in 50 μm squares (i.e. superpixels) for which birthdate time is measured (left). Heatmap of the eight rostro-caudal sections showing the birthdate of superpixels for the EdU Dataset (right). (D) Superpixel birthdate split by supra-structure. (E) Mean birthdate against interquartile range of superpixel birthdate (a measure of the duration of neurogenesis) for supra-structures. (F) Cumulative distribution of superpixel birthdate for supra-structures. (G) Hierarchical clustering of supra-structures based on their mean birthdate and interquartile range reveal a segregation between transient- and sustained-neurogenic regions.

To increase spatial resolution and detect gradients of neurogenesis within structures, we aligned all coronal sections to a reference section at each rostro-caudal level, which allowed us to project labeled cells onto a corresponding anatomical matrix that was subdivided into a grid consisting of 50 x 50 μm superpixels (**Figure 2C**). Defining superpixel birthdate as the average birthdate of all the cells projecting onto it, we generated a total of 31,178 birth-dated superpixels, allowing sub-structural analysis of neurogenesis (**Figure 3D,E**; **Figure S2**). This method, unbiased towards specific structures, mirrored our earlier findings supporting strong ontogenetic bases for anatomical structure formation: superpixels associated with brainstem structures only had early birthdates, whereas forebrain structures had both early and late birthdates (**Figure 2D**). Interestingly, superpixels in nuclear structures such as caudal pontine reticular nucleus (PRNc), external globulus pallidus (GPe) and claustrum (CLA) on average had earlier birthdates than those in laminar structures primary visual area layer 5 (VISp5) and entorhinal area layer 4 (ENTl4) consistent with the dominant presence of nuclear structures in the hindbrain and suggesting distinct temporal constraints over the generation of nuclear vs. laminar stuctures (**Figure S2E**).

**Figure 3.**
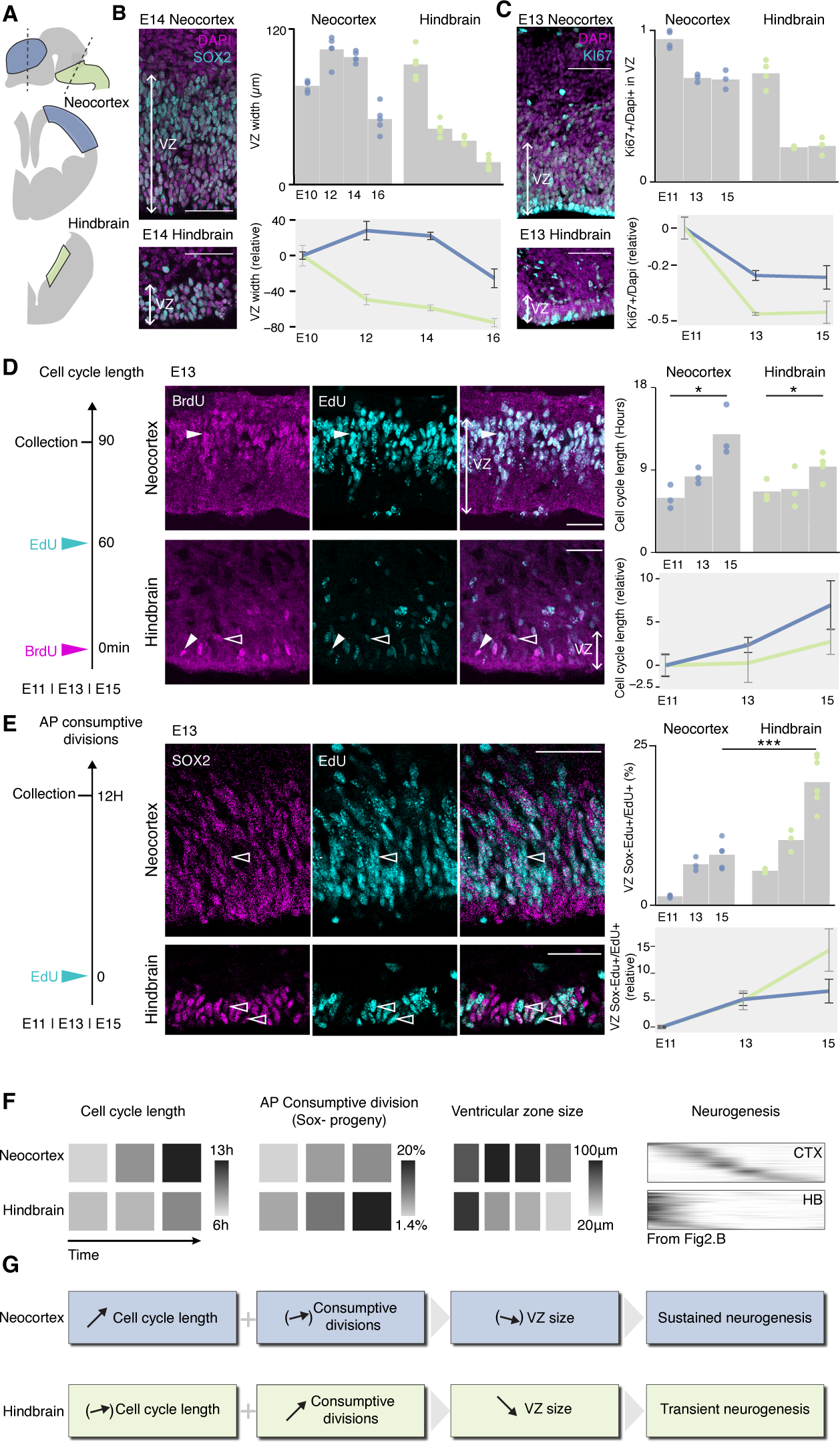
Neocortical and hindbrain apical progenitor have distinct cycling properties and consumption rates. (A) Schematic representation of the VZ regions used in neocortical and hindbrain histological analyses. (B) VZ progenitor marker SOX2 and DAPI staining of E14 neocortex and hindbrain VZs (left); quantification of the width of the VZ (top right); relative temporal evolution to aligned E10 for both regions (bottom right). (C) Proliferative cell marker KI67 and DAPI staining of E13 neocortex and hindbrain VZs (left); quantification of the proportion of KI67^+^ cells over DAPI nuclei in the VZ (top right) and relative temporal evolution to aligned E11 for both regions (bottom right). (D) Double thymidine-analog pulse assay schematic for cell-cycle length measurements (left); representative images (center); quantification (top right) and the relative length aligned to E11 (bottom right). Full arrowhead: BrdU^+^EdU^+^, empty arrowheads: BrdU**^−^**EdU^+^. Schematic representation of the assay used to measure AP-consumptive divisions (left). Representative image (center); quantification of Sox2^−^ EdU^+^ cells over all EdU^+^ cells present in the VZ (top right). Evolution of AP-consumptive divisions across time aligned to E11.5 (bottom right). Arrowheads: SOX2**^−^**EdU^+^ (F) Recapitulative heatmaps of cell-cycle length, consumptive AP division rate and VZ size in the neocortex and the hindbrain along with birthdate-waves of anatomical structures of the cerebral cortex (CTX) and hindbrain (HB), from Figure 2B. (G) Schematic summary of the relationship between cell-cycle length dynamics, AP-consumptive division rates, size of the VZ and neurogenesis type. Scalebars are 100μm. *: *P*<0.05, ***: *P*<0.001.

Using the interquartile range to calculate birthdate dispersion within structures (**Figure 2E**), birthdate distribution analysis over time (**Figure 2F**), and birthdate distribution clustering (**Figure 2G**), we discerned two types of neurogenic dynamics. First, regions in the mid- and hindbrain which demonstrate transient neurogenesis, characterized by rapid cell generation in early embryogenesis without later neurogenesis. Second, regions showing sustained neurogenesis, characterized by cell birth throughout brain development, which we observed primarily in the forebrain and especially the neocortex. Our data, therefore, reveals that rapid, early neurogenesis predominantly occurs in the mid- and hindbrain, while the forebrain and neocortex undergo sustained neurogenesis throughout brain development. These findings beg the question of the cellular mechanisms that lead to prolonged neurogenesis in forebrain structures.

### Neocortical and hindbrain APs have distinct cell-cycle dynamics

The identification of transient and sustained neurogenic regions suggests fundamental differences in the properties and behavior of the progenitor cells from which they arise. Specifically, the tonic generation of neocortical structures may reflect a slower consumption of APs in these regions compared to the hindbrain. To address this possibility, we compared the properties of APs in the neocortex and hindbrain during development (**Figure 3**). As a first approach, we used VZ thickness as a proxy for progenitor numbers, using SOX2 to identify APs and define the limits of this germinal zone between E10 and E16. This strategy revealed that whereas in the hindbrain VZ width decreases rapidly as embryonic days unfold, this decrease is delayed and comparatively limited in the neocortex (**Figure 3A,B**). Using KI67 to identify cycling cells, we also found a progressively decreased fraction of these cells in the hindbrain compared to the forebrain (**Figure 3C**), suggesting differences in cell-cycle kinetics between APs in these two regions. To address this possibility, we measured dynamic changes in AP cell-cycle length and AP-consumptive division rates (i.e., AP divisions giving rise to non-AP cells.) across these two regions during embryogenesis. Using Thymidine analog — EdU and BrdU — double-pulse labeling, we found that the cell-cycle length of neocortical APs increased 2.2-fold between E11.5 and E15.5 (from 6 hours to 12.9 hours respectively, **Figure 3D**), whereas the cell-cycle length of hindbrain APs increased only 1.4-fold during the same period (from 6.7 hours to 9.4 hours; **Figure 3D**). Hence, because of a striking lengthening of their cell cycle, neocortical APs may be less rapidly consumed than their hindbrain counterparts, allowing for prolonged neurogenesis. Supporting this possibility, AP-consumptive divisions were rarer in the neocortex than in the hindbrain, as revealed by a greater fraction of SOX2^−^ cells in the VZ 12h following an EdU pulse injection (**Figure 3E**, 8.1% in neocortex vs 19.8 % in hindbrain at E15.5). Together, these results suggest a scenario in which comparatively slower divisions and fewer consumptive divisions of APs allow prolonged neurogenesis in the forebrain compared to their hindbrain counterparts (**Figure 3F,G**). This led us to ask the question of the molecular processes that could give rise to such spatial and temporal differences in progenitor behavior.

### APs have distinct spatial molecular identities but shared developmental transcriptional programs

To uncover the molecular diversity of APs across space and time, we performed intraventricular injections of FT and collected cells after 1 hour, when juxtaventricular APs are labeled^7,16^. We collected the developing neural tube (excluding the spinal cord) at four different time points — E10, E12, E14, and E16 —, dissociated the tissue, and isolated FT^+^ cells using fluorescence-activated cell sorting (**Figure 4A**). Sorted cells were then processed for single-cell transcriptomics using the 10x Genomics technology (see Methods). To reconstruct the spatial location of progenitors along the neuraxis, we used the Allen Brain Institute’s in situ hybridization (ISH) atlas^17^, which provides developmental gene expression data for 3D spatially-annotated voxels. Using anchor-based data integration, we aligned our dataset with the ISH voxels encompassing the ventricular zone to predict the spatial position of progenitors (**Figure S3A-C**), allowing us to reconstruct spatial and temporal maps of AP gene expression (**Figure 4B**). This approach was validated in an experiment in which the forebrain, midbrain, and hindbrain were microdissected from one another prior to sequencing. In this experiment, both spatial mapping and expression of canonical neural tube segment markers corresponded to the location of the cells (**Figure S3D**).

**Figure 4.**
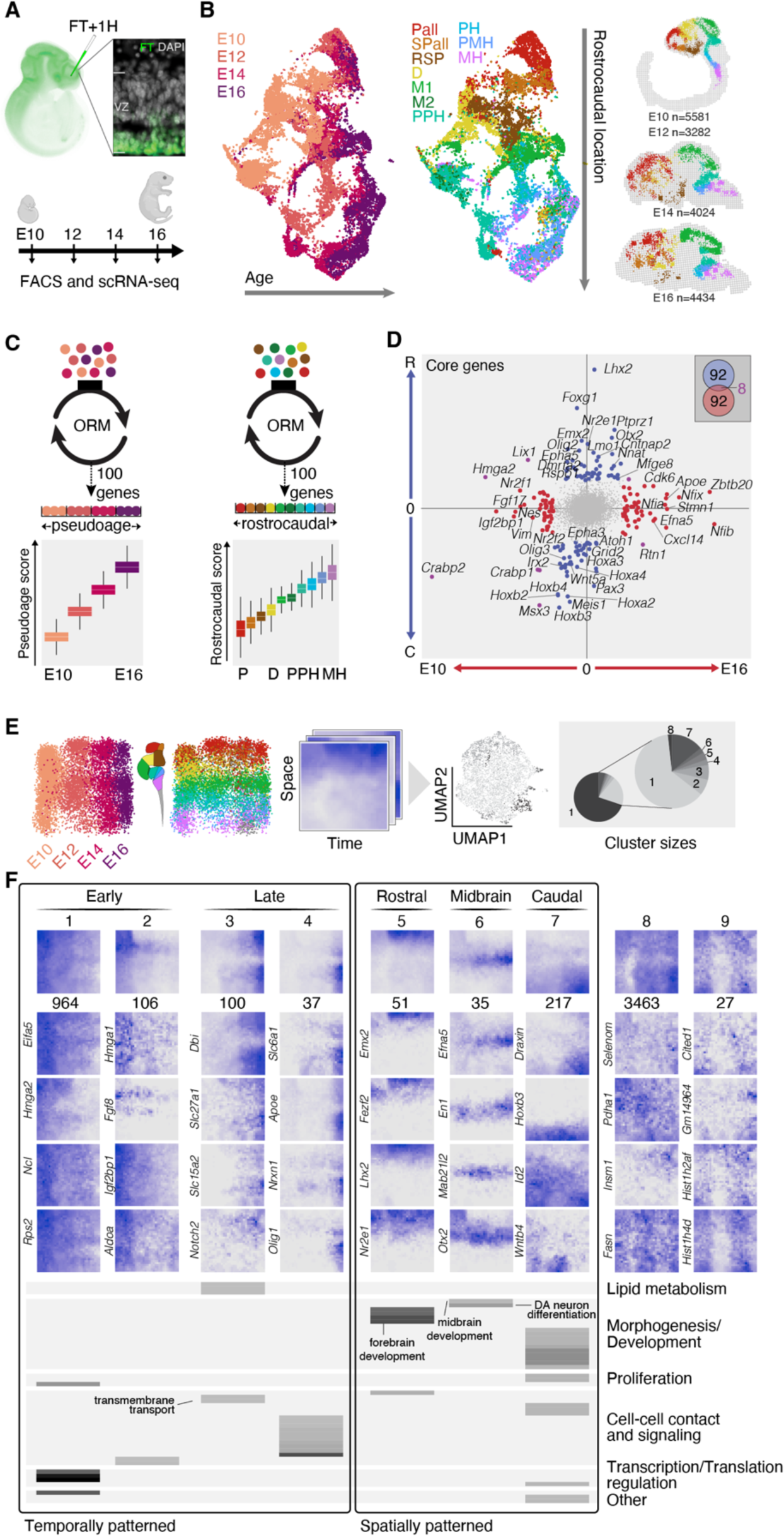
Apical progenitors have distinct spatial molecular identities but shared developmental transcriptional programs. (A) Schematic of AP labeling and single-cell RNA sequencing processing. (B) UMAP representation of AP molecular identity color-coded by age of collection (left). UMAP plot color-coded by rostro-caudal subdivisions of the developing neural tube (center). Result of the spatial mapping of APs onto the developing mouse brain using in situ hybridization voxels of the Allen Brain Atlas with number of cells per age (right), see Figure S3A-C. (C) Schematic of ordinal regression machine learning training to predict pseudo-age and pseudo-rostrocaudal space (top) and cross-validation plots used as pseudo-age and rostro-caudal scores (bottom). (D) Weight of genes from pseudo-age and pseudo-rostrocaudal models with temporal core genes in red, rostro (R)-caudal (C) core genes in blue and common core genes in purple. Venn diagram presenting the distribution of core genes (inset). (E) Generation of gene expression landscapes from pseudo-age and pseudo-rostrocaudal scores (left). UMAP representation of landscapes color-coded by clusters (center); cluster proportions (right). (F) Average landscapes with number of landscapes in each cluster (top) and example genes (middle). Significantly enriched Gene Ontologies for each landscape cluster with either spatial or temporal patterns (bottom).

Analysis of cellular transcriptional identities by Uniform Manifold Approximation and Projection (UMAP) dimensionality reduction revealed that APs were organized based on their embryonic age and rostrocaudal location, with temporal and rostrocaudal identities represented along orthogonal axes (**Figure 4B**). To uncover the molecular programs governing the temporal and spatial properties of APs, we used one ordinal regression machine learning model for each axis, retrieving 100 core genes responsible for the distribution of cells along time and space, respectively (**Figure 4C**). These core genes were essentially specific to the temporal or spatial axis, with only minimal overlap between the two (only 8/200 genes; **Figure 4D**), indicating that regional identity and developmental progression are encoded by different sets of genes. We then combined the two aforementioned models to identify embryonic age- and region-related patterns of AP gene expression. For this purpose, each cell was assigned embryonic age score and rostrocaudal position score. Cells were then embedded within a two-dimensional matrix allowing the display of gene expression profiles as spatio-temporal gene expression maps (**Figure 4E,F**)^18^. To identify archetypical features of gene expression, we performed a UMAP–based analysis of transcriptional maps for all genes, revealing clusters of genes with similar expression dynamics, including genes with clear temporal (**Figure 4F**, Clusters 3-6) or spatial (Clusters 7-9) patterns.

Temporally-patterned genes segregated into “early-on” clusters and “late-on” clusters (**Figure 4F**, Clusters 1-2 and 3-4 respectively). “Early-on” genes (Clusters 1-2) were involved in transcriptional and translational regulation (e.g. the transcription factor *Hmga2* and the translation initiation factor *Eifa5*). This is consistent with the prominence of such processes in early development^18,19^ and in line with the role of protein synthesis/degradation in regulating differences in the timescale of cell divisions between species^20,21^. “Late on” genes (Clusters 3-4) were involved in cell-cell contact and signalling, including for example the cell adhesion protein *Nrxn1*. This is consistent with the increased importance of non-cell-autonomous/exteroceptive processes in regulating properties of late progenitors^18,22^. Genes involved in metabolism such as the peroxisome proliferator-activated receptor *Ppargc1a* (a cell-cycle regulator which acts on cyclins through ATP)^23^ and the transporter *Slc27a1* (a fatty acid transporter which, when inhibited, induces epidermal differentiation)^24^ were also overrepresented in “late on” clusters (Cluster 3), consistent with emerging roles of metabolic states in regulating stem cell properties and developmental tempos^25,26,27^.

Spatially-patterned clusters segregated into rostrally-enriched genes (Cluster 5), midbrain-enriched genes (Cluster 6) and caudally-enriched genes (Cluster 7). These included classical rostro-caudal neural tube patterning transcription factors such as *Lhx2*, *Emx2* and *Hoxb3*^5,28,29^. Also *En1, En2* and *Otx2*, which control dopamine neuron differentiation were prominently observed in APs of the midbrain — the area where these neurons originate (Cluster 6)^30,31^.

Finally, most genes were expressed in diffuse or focal patterns (Clusters 8-9, containing 70% of all genes). These included for example *Insm1* (late on and rostral), which promotes the generation of intermediate progenitors^32^, consistent with the regional differences in indirect neurogenesis identified earlier (see **Figure S2**)^14,33^. Metabolism-related and mitochondrial genes were also expressed within these clusters, including the peroxisome proliferator activated receptors *Ppard*^34^ and fatty acid synthase *Fasn*^35^, together further supporting the functional relevance of the transcriptional programs identified here.

### The mitochondrial protein *Fam210b* regulates region-specific AP cycling properties during development of the forebrain and hindbrain

The findings above suggest that the properties of APs along the neural tube are molecularly encoded by distinct spatio-temporally-regulated gene clusters. If proven functional, this transcriptional organization might be the key to the different cell-cycle dynamics in rostral (i.e. neocortical) and caudal (i.e. hindbrain) APs.

To pinpoint gene candidates that could explain the extended cell cycling of neocortical APs, we built a synthetic landscape that matched this cellular feature in space and time (i.e. high late-forebrain expression) and looked for closest neighbor genes on the UMAP space that could serve critical functions (**Figure 5A**). A key candidate that emerged was a mitochondrial inner membrane protein known to regulate erythroid progenitor divisions^36^, FAM210B, whose mRNA was expressed where and when AP cell cycling is slowest (**Figure 5A**). If this late-onset gene was suppressed in the neocortex, it could thus conceivably give rise to hindbrain-like AP cycling properties (as per **Figure 3G**).

**Figure 5.**
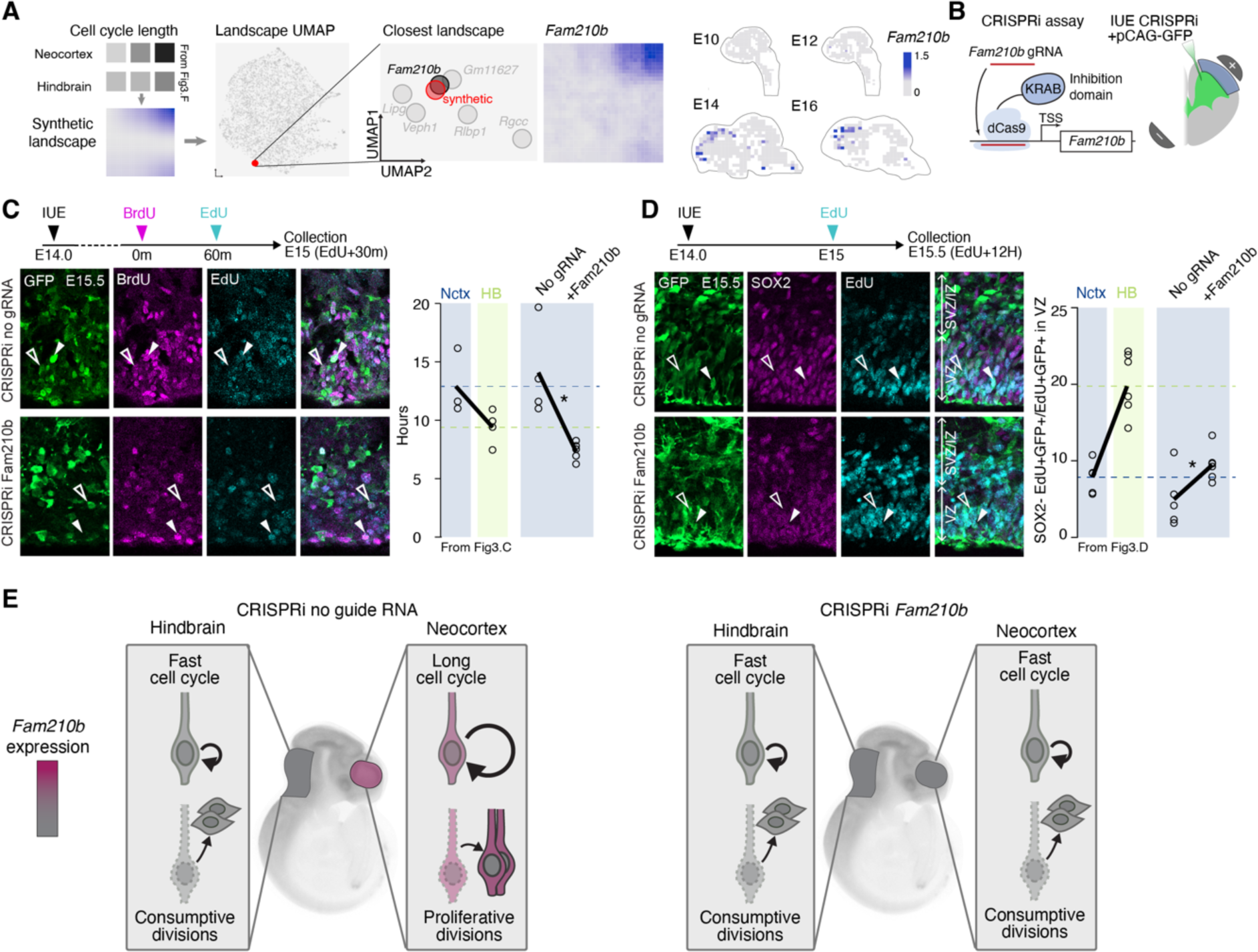
Repression of the mitochondrial protein Fam210b reduces cell-cycle length and increases consumptive divisions in the developing neocortex. (A) Generation of a synthetic landscape based on cell-cycle length measurements (left, from Figure 3), UMAP plot with synthetic landscape position highlighted, and zoom-in of the synthetic landscape’s neighborhood (middle). Fam210b landscape and spatial distribution (right). (B) Schematic of the CRISPRi assay (left) and in utero electroporation targeting strategy (right). (C) Measurement of cell-cycle length with dual thymidine analog pulse-chase label following IUE of CRISPRi system (top left). Representative image of electroporated cells with GFP, BrdU and EdU stainings (bottom left) and effects on cell-cycle length (neocortex and hindbrain data reported from Figure 3C; right). Empty arrowheads: GFP^+^BrdU^+^EdU**^−^**, full arrowheads: GFP^+^BrdU^+^EdU^+^ (D) Schematic of consumptive division rates measurement following IUE of CRISPRi strategy (top left) with representative images of GFP, Sox2 and EdU (bottom left) and effects on AP-consumptive divisions (neocortex and hindbrain data reported from Figure 3D; right).. Empty arrowheads: GFP^+^SOX2**^−^**EdU^+^, full arrowheads: GFP^+^SOX2^+^EdU^+^. (E) Schematic summary of the results. *: P < 0.05.

To investigate this possibility, we used CRISPR inhibition (CRISPRi) to repress *Fam210b* transcription. We first delivered the CRISPR system and guide into neocortical AP at E14.5 through *in utero* electroporation (IUE), thereby confining its repression to the forebrain (**Figure 5B**). Next, we measured cell-cycle length and AP consumptive division rates in CRISPRi-*Fam210b* cells, as previously conducted to examine region-specific cell-cycle characteristics (see **Figure 3**). Confirming our hypothesis, the results showed that repression of *Fam210b* expression decreased cell-cycle length to the levels usually observed in the hindbrain (**Figure 5C**) and increased AP consumptive divisions, as demonstrated by an increase in the production of SOX2^−^ cells (**Figure 5D**). Together, these findings underscore the role of region-specific cell-cycle regulation by spatially and temporally distributed molecular mechanisms in controlling AP consumption timing and, by extension, regional brain generation (**Figure 5E**).

## DISCUSSION

Our results provide a parsimonious mechanism to regulate neuronal production in different parts of the brain. In regions that expand during development and phylogenesis, such as the neocortex, neurogenesis is sustained across brain development, while it is confined to the first few days of brain formation in hindbrain regions that develop comparatively less. At the cellular level, sustained neocortical neurogenesis is made possible by a progressive and marked lengthening of the cell cycle and lower levels of AP-consumptive divisions, which together preserve the progenitor pool for longer periods. Repression of a forebrain-specific gene, *Fam210b*, provides a proof-of-principle demonstration that spatially-restricted expression of a specific gene is sufficient to confer region-specific properties to APs along the neuraxis. Hence the time window during which neurons are generated in each brain region appears to be a critical determinant of this structure’s expansion during development and, arguably, evolution.

The mechanisms that regulate the timing of cellular processes across species have recently begun to be addressed^37,38,39^. For example, the cellular segmentation clock is directly influenced by the kinetics of HES7 – inhibiting or promoting HES7 protein synthesis respectively slows down or speeds up segmentation – and differences in the timescale of this clock across species reflect differences in the stability of this protein^20^. Similarly, differences in protein stability have also been involved in regulating the differences in the cell-cycle duration of mouse and human embryonic stem cells *in vitro*^21^. More recently, the pace of mitochondrial development and metabolic activity has been shown to set the pace of neuronal development *in vitro* and *in vivo*^26^. Supporting a central role for metabolism in driving cell cycle properties, early in corticogenesis, fast-cycling, proliferative progenitors prioritize anaerobic glycolysis while (mitochondria-dependent) oxidative processes emerge at later stages, in conjunction with the emergence of consumptive divisions^27^.

Our findings reveal that within the developing embryonic brain, cellular clocks do not tick at the same pace everywhere, as cell-cycle lengthening and proliferative divisions occur at a more protracted pace in the forebrain than in the hindbrain. Remarkably, we find that the mitochondrial protein FAM210B, which also regulates erythroid progenitor function^36^, is involved in regulating cell-cycle duration in the forebrain, consistent with the role of metabolic activity in regulating developmental tempo discussed above. It will thus be interesting, in future studies, to compare metabolic rates and activities across brain regions as these may participate in conferring region-specific properties to progenitors and their daughter cells.

By comparing data obtained with the EdU and the FT birthdating datasets, we were able to examine the extent of indirect neurogenesis in regions outside of the neocortex. Our results confirm and extend previous findings^13,14^ by showing extensive indirect neurogenesis in the thalamus and, to a lesser degree, in the striatum. Hence, while indirect neurogenesis is believed to significantly contribute to the expansion of the neocortex in mammals^40^, the thalamus, to which it is intimately functionally tied, may also experience substantial indirect neurogenesis in various species.

Given the vital autonomous functions subserved by hindbrain structures, there must have been a strong selection pressure for them to be generated rapidly. On the other hand, extended neurogenesis in the forebrain may facilitate interactions with environmental factors or activity-related processes, potentially adapting neurons and circuits to particular environmental conditions. Supporting this possibility, we have shown that forebrain APs become more exteroceptive during corticogenesis, including through hyperpolarization^18,22^, which may allow thalamocortical afferents to modulate progenitor behavior^41,42^.

Anatomical correspondences across developing species have long struck embryologists, leading to Haeckel’s now largely outdated theory that “ontogeny recapitulates phylogeny”^43^ – it is understood that newer structures are not simply added onto older ones. Instead, as we show here, both forebrain and hindbrain structures develop initially simultaneously, with the difference that forebrain structures have acquired the ability to undergo sustained neurogenesis. It will be interesting, in future research, to examine to which extent region-specific properties of progenitors are cell-intrinsically determined and exhibit plasticity in various contexts.

## ACKNOWLEDGEMENTS

We thank the Genomics Platform, Bioimaging Facility and FACS Facility of the University of Geneva; Nicolas Liaudet for expertise on image analysis; A. Benoit for technical assistance; J. Prados for assistance with bioinformatics analyses; Q. Lo Giudice and the Informatics service of the faculty of medicine of the University of Geneva for support in publishing the website, L. Frangeul for assistance in select early experiments, and all members of the Jabaudon laboratory for their comments on the manuscript as well as members of the Tole laboratory for constructive exchanges during the project. The Jabaudon laboratory is supported by the Swiss National Science Foundation, the Carigest Foundation, the Société Académique de Genève FOREMANE Fund, and the European Research Council. Robin J. Wagener was supported by the Deutsche Forschungsgemeinschaft (DFG) Research Fellowship Wa 3783/1.

## AUTHOR CONTRIBUTIONS

Tissue collection, staining and image acquisition for the birthdating atlas were performed by RW. Delineations of birthdating atlas sections were done by RW, NB, PA, ALa, ALo, EM and SF. Webtool design, implementation and data extraction for the birthdating atlas were done by NB. Single-cell transcriptomics experiments were performed and analyzed by NB. The embryonic experiments, histology, and analyses to address the cell-cycle properties of progenitors were performed by AJ, NB and DF. The CRISPRi design, experiments and analyses were performed by AJ. Figure design and execution were done by NB with the help of DJ and AJ. Manuscript redaction was done by DJ and NB and all authors reviewed the manuscript. DJ, NB, RW and AJ designed the experiments.

## METHODS

### Declaration of generative AI and AI-assisted technologies in the writing process

During the preparation of this work the authors used Chat-GPT in order to streamline some parts of the text. After using this tool, the authors reviewed and edited the content as needed and take full responsibility for the content of the publication.

## RESOURCE AVAILABILITY

### Lead contact

Further information and requests for reagents and recourses should be directed to D. Jabaudon.

### Material availability

This study did not generate new material or reagents.

### Data and code availability

- Single-cell RNA sequencing data will be deposited on GEO and be publicly available as of the date of publication. Accession number will be listed in the key resource table.
- This paper does not report original code.

## EXPERIMENTAL MODEL AND STUDY PARTICIPANT DETAILS Mice

The experiments were conducted following Swiss laws and received approval from the Geneva Cantonal Veterinary Authorities and its ethics committee. The study adhered to the ARRIVE guidelines. CD1 male and female mice from Charles River Laboratory were used. Matings were conducted overnight for the birthdating atlas, with the subsequent morning designated as time E0.5. Mice for embryonic assays and histology were mated within a 3-hour timeframe, which was designated as E0.

## METHOD DETAILS

### EdU injections (Dataset 1)

150μl of EdU (1mg/g) was injected intraperitoneally in pregnant dams at the indicated gestation time points.

### *In utero* FlashTag injections (Dataset 2)

Pregnant dams were administered 2-3% isoflurane anesthesia at specific gestation time points and placed on a warm operating table. Small incisions were made in the abdomen to expose the uterine horns, and FlashTag (CellTrace CFSE) was injected into the ventricles as previously described^7^. A volume of 414 nL of FlashTag was injected into the lateral ventricle. After the procedure, the uterine horns were returned to the abdominal cavity, and the peritoneum and skin were independently sutured. The mice were placed on a heating pad until they recovered from anesthesia.

### Chronic BrdU delivery

For chronic BrdU delivery, osmotic pumps loaded with 16 mg/mL BrdU 1:1 in PBS and DMSO were utilized. To cover the required delivery period, 0.1 μL per hour osmotic pumps were used, specifically the 1003D Alzet pump for a duration of three days and the 2001 Alzet pump for a duration of seven days. The osmotic pumps were introduced into the peritoneal cavity during *in utero* injections.

### Post mortem tissue collection

E10 and E11 embryos were decapitated whereas for all other embryos, the entire brain was dissected out of the head and immersed in 4% PFA/PBS overnight. The next day, the samples were washed with 1X PBS and transferred into 20% Sucrose/PBS overnight. After freezing the tissue in OCT, cryosectioning was performed at a thickness of 20μm. Slides were kept at −80C before using for immunostaining.

Postnatal brains were collected by conducting intracardiac perfusion of 10% sucrose followed by 4% paraformaldehyde (PFA) while the mice were under thiopental anesthesia. Subsequently, the postnatal brains were fixed overnight at 4°C in 4% PFA. The brains then underwent equilibration in a 20% sucrose PBS solution before being embedded in OCT medium. Using a Leica cryostat, the brains were sliced into 60 μm coronal sections. These sections were then maintained in a free-floating arrangement.

### *In utero* electroporations

Electroporations were performed as previously described with a few modifications^22^. Plasmid DNA was injected at a total concentration of 2μg/μL in the lateral ventricles of E14 embryos and electroporation was performed with 42V.

### CRISPRi gRNA design and cloning

CRISPick^44^ was used to design gRNA targeting Fam210b for SpyoCas9 CRISPRi using reported scoring model^45^. Top candidate gRNA sequence was ordered as single stranded oligo from IDT containing backbone homology arms for cloning (AAAGGACGAAACACC**CAGCGTCAACAGCCCGGCCA**GTTTTAGAGCTAGAA, gRNA sequence highlighted in bold). This sequence was cloned into pX458-Ef1a-dCas9-KRAB-MECP2-H2B-GFP-NogRNA backbone by first digesting using BbsI and then performing NEBuilder HiFi DNA assembly reaction.

#### Plasmids

Injected plasmid were: pCAG-GFP (0.5μg/μl), pX458-Ef1a-dCas9-KRAB-MECP2-H2B-GFP-NogRNA (1.5μg/μl) or pX458-Ef1a-dCas9-KRAB-MECP2-H2B-GFP-Fam210b-gRNA (1.5μg/μl).

#### Intraperitoneal injections

Injection were performed as previously described with a few modifications^22^. EdU (7.5mg/ml) or BrdU (10mg/ml) were dissolved in 0.9% NaCl and injected volumes corresponded to 50mg/kg of animal weight.

### Immunohistochemistry

#### Birthdating dataset

All free-floating sections were washed three times 10 min in PBS at room temperature. BrdU-containing sections were denaturated prior to blocking by incubating them in 2 N HCl at 37°C for 30 min and washed twice in PBS for 30 min. The sections were then incubated 1h at room temperature in blocking solution (10% normal horse serum, 0.1% Triton-X 100 in PBS) and then incubated at 4°C with primary antibody diluted in the same blocking solution. Sections were then washed four times in PBS for 15 min and incubated 2h with corresponding secondary antibodies diluted in blocking solution. After washing again 3 × 15 min with PBS, sections were mounted in Sigma Fluoro-mount (#F4680). For EdU sections, the Click-it chemistry was used following the manufacturer’s instructions (Invitrogen). Sections were then incubated with DAPI (1:1000 in PBS) for 10 minutes at RT and washed in PBS for 10 minutes.

#### Embryonic histology

For EdU click-it revelation, a custom protocol composed of mixing 1X PBS (429ul) with 100mM of CuSO4_4_ (20ul, 4mM final), AF azide 647 (1ul) and 1M of Sodium L-Ascorbate (50ul, 100mM final, used either fresh or stored at −20C) was used. Components were mixed in the above listed order and immediately added on the slides for 30mins. This was done at the end of the immunostaining.

For pulse-chase experiments, to limit the cross reactivity of the BrdU antibody with EdU, a blocking protocol was used as previously described with minor modifications^46^. Before immunostaining, OCT was first washed from slides by immersing in 1X PBS. Then slides were treated with 1X PBS + 0.5% Triton for 10 min. After permeabilization, click-it reaction was performed as described above with a fluorescent azide. After the first click-it reaction, slides were washed three times with 1X PBS and then the same click-it solution was prepared but by substituting AF azide 647 with a non-flourescent Azidomethyl phenyl sulfide (20mM final). Slides were treated with this solution for 30mins and washed with 1X PBS three times. A final click-it reaction was performed without azide for 10 min. Finally, the slides were then immersed in a solution of 20mM EDTA/PBS for 30 min. The slides were then washed in 1X PBS three times and incubated with primary antibodies overnight at RT in 1X Exonuclease buffer containing 0.1U/μl of Exonuclease III. The secondary antibodies were added the next day for 2h and sections were processed with same protocol as the postnatal tissue.

### Imaging

Birthdating assay were imaged with Epifluorescence ZEISS Axio Scan.Z1 slide scanner. Embryonic histology images were obtained using either a Leica Stellaris confocal mounted with 20x/0.75 objective and ZEISS LSM 800 mounted with 20 × 0.5 CFI Plan Fluor WD:2.1 mm or 40 × 1.3 CFI Plan Fluor DIC WD:0.2 mm objectives.

### Single-cell RNA-seq capture

Single-cell library captures were performed on single embryos and the dataset is composed of 2 libraries for E10, and 3 libraries for E12, E14 and E16 time points.

#### Cell dissociation and FAC-sorting

Pregnant females were sacrificed one hour after performing *in utero* FlashTag (FT) injections. The neural tubes of the embryos were collected in ice-cold Hank’s Balanced Salt Solution (HBSS), and the forebrain, midbrain, and hindbrain segments were microdissected under a stereomicroscope. The tissue was then digested using TrypLE (Gibco) at 37°C for three minutes, followed by the addition of FACS buffer (which consisted of 2mg/mL glucose, 0.1% BSA, 1:50 Citrate Phosphate Dextrose from Sigma (C7165), 10U/mL DNase I, and 1μM MgCl2). Mechanical dissociation was carried out by pipetting up and down. The resulting cell solution was passed through a 70 μm cell strainer and centrifuged at 150 G for 5 minutes. After resuspending the cells in FACS buffer, as previously described in the reference, the 10% brightest FlashTag-labeled cells were sorted using a MoFloAstrios device from Beckman. 10,000 cells were sorted for each of the three primary segments (forebrain, midbrain, and hindbrain) before subsequently pooling them back together for scRNA-seq capture. An exception was made for one E12 collection which scRNA-seq capture was done individually for forebrain, midbrain and hindbrain (**Figure S3D**).

#### Single-cell RNA sequencing

42 μL of cell suspension was captured using 10X Genomics Chromium Single Cell 3’ v3 reagents and protocol. The quality control of the cDNA and libraries was performed using Agilent’s 2100 Bioanalyzer. Subsequently, the libraries were sequenced using the HiSeq 2500 sequencer. The resulting FASTQ files obtained from the sequencing were processed and mapped with 10X Genomics Cell Ranger pipeline (version 3.0.2) using the GRCm38 mouse genome as a reference.

## QUANTIFICATION AND STATISTICAL ANALYSIS

### Image analysis and quantification

#### Cell detection of birthdating datasets

Images of the birthdating Atlas were processed for cell detection on FT or EdU channel using MetaXpress software (v.5.1.0.41, Molecular Devices) using minimal intensity requirements. Clustering analysis was used on detected cells intensity in order to detect potential thresholds for cell filtering. The intensity threshold was then selected manually on unlabeled images to avoid bias (**Figure S1C**). For the FT and chronic BrdU dataset, a set of images were taken with confocal microscopy and FT+BrdU-cells were manually annotated. These annotations were used to train a machine learning logistic regression to exclude FT^+^BrdU^+^ from FT^+^BrdU^-^ cells. In order to avoid false positive detection, all pial surfaces all sections were manually removed.

#### Birthdating images alignment to reference section

All FT and EdU images were aligned to a reference section for each rostro-caudal level using Matlab. One reference section was chosen among the 48 sections of the dataset for each level (8 ages per 3 replicated per 2 datasets) and pairs of points were placed as anatomical landmarks on VGLUT2 staining between the reference and the section to align. The image was then aligned to the reference using the fitgeotrans function and local weighted mean (lwm) transformation. The resulting matrix of transformation was used to project detected cell positions into the reference section and thus align all ages and replicates on the same 2D space (**Figure 2C**).

#### Birthdate-wave generation and analysis

Existing adult anatomy annotation tools (specifically, QuickNII and the Wholebrain R package) did not perform well on P7 sections, such that the anatomical structures of all 384 brain sections were manually annotated based on the adult mouse Allen Brain Atlas (Coronal sections, http://mouse.brain-map.org/). EdU and FT positive cells were counted per anatomical structure and normalized by the area of the region to get density values for each anatomical structure at each age and each replicate. A loess smoothing was used on density values for all the replicates across ages to generate a 71 point-based birthdate wave. All waves were then normalized to the same area under the curve (AUC=1). The clustering of Figure 2G was perfomed using *hclust* function in R. The distance measured prior to clustering is an euclidean distance between 500 randomly sampled superpixels from each supra-structure.

#### Embryonic image analysis

Images of KI67 staining (**Figure 3C**) were analyzed using a semi-automated cell detection pipeline in Matlab. In order to detect nuclei, an intensity threshold was applied on DAPI staining images, followed by a watershed algorithm with the *watershed* function in Matlab. The resulting detected spots were filtered in order to remove spots that were too small or too large. The intensity of KI67 staining was average for each of the detected nuclei and manual thresholding was done to determine KI67^+^ cells.

For consumptive division assay, three 100 μm areas on multiple sections were imaged per animal per stage per region of the brain. An imageJ plugin as previously implemented^47^ was used to count EdU^+^ cells that first analyses particles (min=4 and max=15) in a defined region of interest, sets a user-defined threshold for positive signal and then utilizes watershedding to segment detected particles into individual cells. Once all EdU cells were counted, SOX2 was used to define EdU^+^SOX2^+^ or EdU^+^SOX2^−^ cells in the ventricular zone. For CRIPSRi experiment, only GFP^+^ cells in the image were counted.

For pulse chase experiments, EdU^+^ cells were counted using the above-mentioned plugin with SOX2 labelling as an identifier of VZ, then BrdU^+^ cells were analyzed by counting manually. Formula listed in this study^48^ was used for quantifying cell-cycle length. S-phase = (SOX2^+^BrdU^+^EdU^+^ / SOX2^+^BrdU^+^EdU^−^); total cell cycle = S-phase/(BrdU^+^SOX2^+^/SOX2^+^). For CRIPSRi experiment, only GFP^+^ cells in the image were counted. All other image quantifications were performed manually using standard Fiji functionalities.

Quantification shown in **Figure 3** and **Figure 5** are mean ± SD. Welch Two Sample t-test were performed and statistical indication follow the convention: *:*P* < 0.05, **:*P* < 0.01, ***:*P* > 0.001. In **Figure 3** and **5** each dot represents data from one embryo. **Figure 3B** Neocortex and Hindbrain n = 5 for all stages. **Figure 3C** E11.5 Neocortex and Hindbrain n = 4, for all other stages n = 3. **Figure 3D** Neocortex E11.5, E13.5 and E15.5 n = 3; Hindbrain E11.5 and E13.5 n = 3, E15.5 n = 4. Test between E11.5 and E15.5 Neocortex: *P*=0.017; Test between E11.5 and E15.5 Hindbrain *P* = 0.02. **Figure 3E** Neocortex E11.5 n=6, E13.5 n=3, E15.5 n = 5; Hindbrain E11.5 n = 3, E13.5 n = 3. E15.5 n = 7; Test between E15.5 Neocortex and E15.5 Hindbrain: *P* = 3.46e-5. **Figure 5C** no gRNA n = 4, *+Fam210b* n = 5; *P* = 0.021. Figure 5D no gRNA n = 5, *+Fam210b* n = 6; *P* = 0.027.

### Single-cell RNA-seq Data Analysis

All single-cell transcriptomics analyses were performed using R Statistical Software (v4.2.2), the Seurat package and Bioconductor packages. Graphs and visualization were generated with the *ggplot2* package.

#### Quality control

All single-cell transcriptomics analyses were performed using R Statistical Software (v4.2.2), the Seurat package and Bioconductor packages.

Single libraries were processed for quality control as Seurat objects. All cells with less that 1,000 detected genes, less than 5,000 detected RNA molecules and more than 10% of mitochondrial RNA expression were removed. In addition, all genes that were not expressed in more that 5% of cells were removed from the analysis. The doublet finder package was used to detect and exclude doublets from the dataset. In addition, a score of neuronal and intermediate progenitors identity was measured as the mean log2 normalized UMIs per 10,000, using published scRNA-seq markers of these types^18^. All cell showing a neuron score > 0.15 and an IP score > 0.6 was discarded of further analysis.

#### Spatial mapping

ISH voxel data for E11.5, E13.5 and E15.5 mouse brains were downloaded from the Allen Brain Institute API (http://api.brain-map.org/) and processed as Seurat objects.

The dataset from each age were independently aligned on the voxels using the integration pipeline provided with Seurat (v3. and further)^49^. Cells from E10 and E12 time points were aligned to the E11.5 voxels while cells from E14 were aligned on E13.5 voxels and cells from E16 to E15.5 voxels. Integrated cells and voxels were then projected on PCA and UMAP space. Each cell was assigned the position and considered as part of the anatomical area of the closest voxel on the UMAP projection.

#### Pseudo-age and pseudo-rostrocaudal reconstruction

Pseudo-age and pseudo-rostrocaudal scores were reconstructed using ordinal regression models were performed with the bmrm package as previously described^18^. Briefly, the regularized ordinal regressions were used to separate embryonic age of collection (E10, E12, E14, E16) and rostro-caudal positioning (Pall, Spall, RSP, D, M1, M2, PPH, PH, PMH, PM). During modeling, each gene is assigned a weight according to its importance in predicting cells in the correct order. The most 50 negatively and most 50 positively weighted genes are retrieved as core gene of the process being predicted (Figure 4D). In order to avoid overfitting, a cross-validation was perfomed for each model; Iteratively, 10% of cells were removed and the model was trained on 90% remaining cells. The resulting model was used to predict the 10% of cells that was not include in the training. This processed was then done 10 times until each cell had a prediction value. These cross-validation values are used as pseudo-age and pseudo-rostrocaudal scores (Figure 4C).

#### Spatio-temporal gene expression landscapes

Procedure of landscape generation was performed as previously described^18^. Cells were arranged in a two-dimensional grid based on the pseudo-age and pseudo-rostrocaudal score using *tanh* function to align the cell on a square grid. The expression of the top 5000 variable genes was measured at each point of the 32 x 32 grid by averaging the expression of its nearest 25 neighboring cells. All expression maps were scaled to the sum of all their values prior to principal component analysis. K-nearest neighbors clustering was performed (K = 20) on the 8 principal components generated and a 2D UMAP space was computed for visualization. The Gene Ontology enrichment analysis was performed with the ClusterProfiler R package on biological processes, p-value were adjusted with Benjamini-Hochberg procedure (**Figure 5F**).

## ADDITIONAL RESOURCES

All birthdating images and waves are available on our webtool https://neurobirth.org

KEY RESOURCE TABLE.

KEY RESOURCE TABLE
| REAGENT or RESOURCE | SOURCE | IDENTIFIER |
| --- | --- | --- |
| **Antibodies** |  |  |
| Rat monoclonal anti-BrdU (1 :200) | Abcam | Cat# AB6326 RRID:AB_305426 |
| Rabbit polyclonal anti-Fluorescein (1 :1000) | Abcam | Cat# AB19491, RRID:AB_444949 |
| Guinea pig polyclonal anti-VGLUT2 (1:5000) | Millipore | Cat# AB2251-I, RRID:AB_2665454 |
| Rabbit anti-KI67 (1:250) | Abcam | Cat# AB15580 RRID:AB_443209 |
| **Chemicals, peptides, and recombinant proteins** |  |  |
| BrdU (5-Bromo-2'-deoxyuridine) | Thermofisher | Cat# H27260 |
| Click-iT EdU Alexa Fluor 647 | Thermofisher | Cat# C10419 |
| EdU | Thermofisher | Cat # A10044 |
| CellTrace CFSE Cell Proliferation Kit | Thermofisher | Cat# C34554 |
| BbsI | NEB | Cat# R0539S |
| NEBuilder HiFi DNA | NEB | Cat# E2621S |
| AF azide 647 | Lumiprobe | Cat# A6830 |
| Azidomethyl phenyl sulfide | Sant Cruz Biotechnology | sc-233899 |
| Chromium Single Cell 3' GEM Kit v3 | 10x Genomics | PN-1000077 |
| Chromium Single Cell 3' Library Kit v3 | 10x Genomics | PN-1000078 |
| Chromium Single Cell 3' Gel Bead Kit v3 | 10x Genomics | PN-1000076 |
| Chromium Chip B Single Cell Kit | 10x Genomics | PN-1000073 |
| Exonuclease III | Thermofisher | Cat# EN0191 |
| **Experimental models: Organisms/strains** |  |  |
| CD-1 mouse | Charles River | RRID:MGI:5649524 |
| **Oligonucleotides** |  |  |
| pX458-Ef1a-dCas9-KRAB-MECP2-H2B-GFP | Addgene | Car#175573<br>RRID:Addgene_175573 |
| **Software and algorithms** |  |  |
| Adobe Illustrator CC | Adobe | RRID:SCR_010279 |
| Adobe Photoshop CC | Adobe | RRID: SCR_014199 |
| ImageJ/FIJI | ImageJ | https://imagej.net |
| R Project for Statistical Computing | R-project | RRID:SCR_001905<br>http://www.r-project.org/ |
| MetaXpress software | Molecular Devices | RRID:SCR_016654<br>https://www.moleculardevices.com/products/cellular-imaging-systems/acquisition-and-analysis-software/metaxpress |
| MATLAB | Mathworks | RRID:SCR_001622<br>http://www.mathworks.com/products/matlab/ |
| Other |  |  |
| Alzet pump | Charles River | ALZT1003D |
| Alzet pump | Charles River | ALZT2001 |

## SUPPLEMENTAL FIGURES

**Figure S1.**
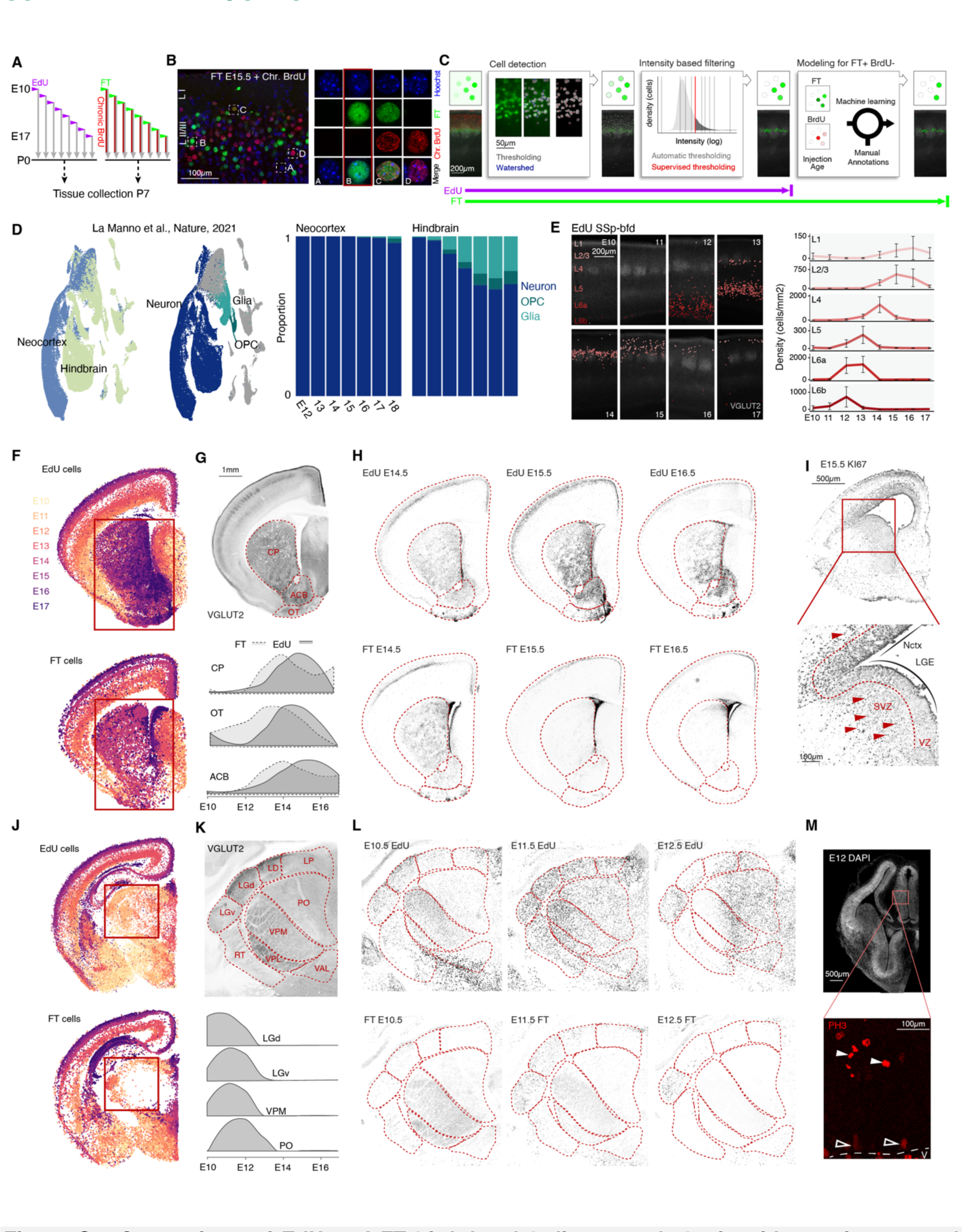
Comparison of EdU and FT birthdate-labeling reveals brain-wide spatio-temporal patterns of direct vs. indirect neurogenesis. (A) Schematic representation of the birthdating strategies using an EdU pulse (Dataset 1) or a FT pulse and chronic BrdU (Dataset 2). (B) P7 image of E15.5 FT and chronic BrdU neocortical section (left) and example of staining combinations present, FT^+^BrdU^−^ cells represent directly VZ-born neurons (highlighted, left) See Govindan et.al^7^. (C) Schematic representation of the cell detection and filtering pipeline consisting in cell detection, intensity-based filtering and prediction of FT^+^BrdU^−^ cells from a machine learning model trained on confocal stacks. Steps of the image analysis applied to EdU and FT datasets are shown (bottom). (D) UMAP plot of neocortical (dorsal pallium) and hindbrain single-cell RNA-seq from the dataset of La Manno *et.al* ^11^. Proportion of cell type for neurons, OPCs and glia normalized to 1 per age. (E) Somatosensory barrel field VGLUT2 images with reconstructed EdU^+^ cells color-coded by their cortical layer location (left) and cell densities of corresponding cortical layers; error bars correspond to standard deviation (top right). (F,J) Projection of all detected EdU^+^ and FT^+^ cells onto a common space color-coded by birthdate. (G) VGLUT2 staining of P7 mouse brain section with striatum nuclei highlighted (top) and their corresponding birthdate-wave for EdU and FT (bottom). (H) EdU (top) and FT (bottom) stainings of striatal region at E14.5, E15.5 and E16.5. (I) Proliferation marker KI67 in E15.5 forebrain (top) and high magnification of lateral ganglionic eminence and neocortex. Arrowheads: KI67^+^ cells in the SVZ. (K) VGLUT2 staining of P7 mouse brain section with thalamic nuclei highlighted (top) and their corresponding birthdate-wave for EdU and FT (bottom). (L) EdU (top) and FT (bottom) stainings of caudal thalamic area at E10.5, E11.5, and E12.5. (M) Low-magnification of DAPI staining of E12 mouse brain coronal section (top) and highlight of mitotic marker PH3 in the prosomere 2 domain (full arrows: PH3^+^ cells in the SVZ; empty arrows: PH3^+^ cells in the VZ). ACB: Nucleus accumbens, CP: Caudoputamen; LD; Lateral dorsal nucleus of thalamus, LGd: Dorsal part of the lateral geniculate complex; Ventral part of the lateral geniculate complex; LP: Lateral posterior nucleus of the thalamus; OT: Olfactory tubercule; PO: Posterior complex of the thalamus; VAL: Ventral anterior-lateral complex of the thalamus; VPL: Ventral posterolateral nucleus of the thalamus; VPM: Ventral posteromedial nucleus of the thalamus; V: ventricular wall.

**Figure S2.**
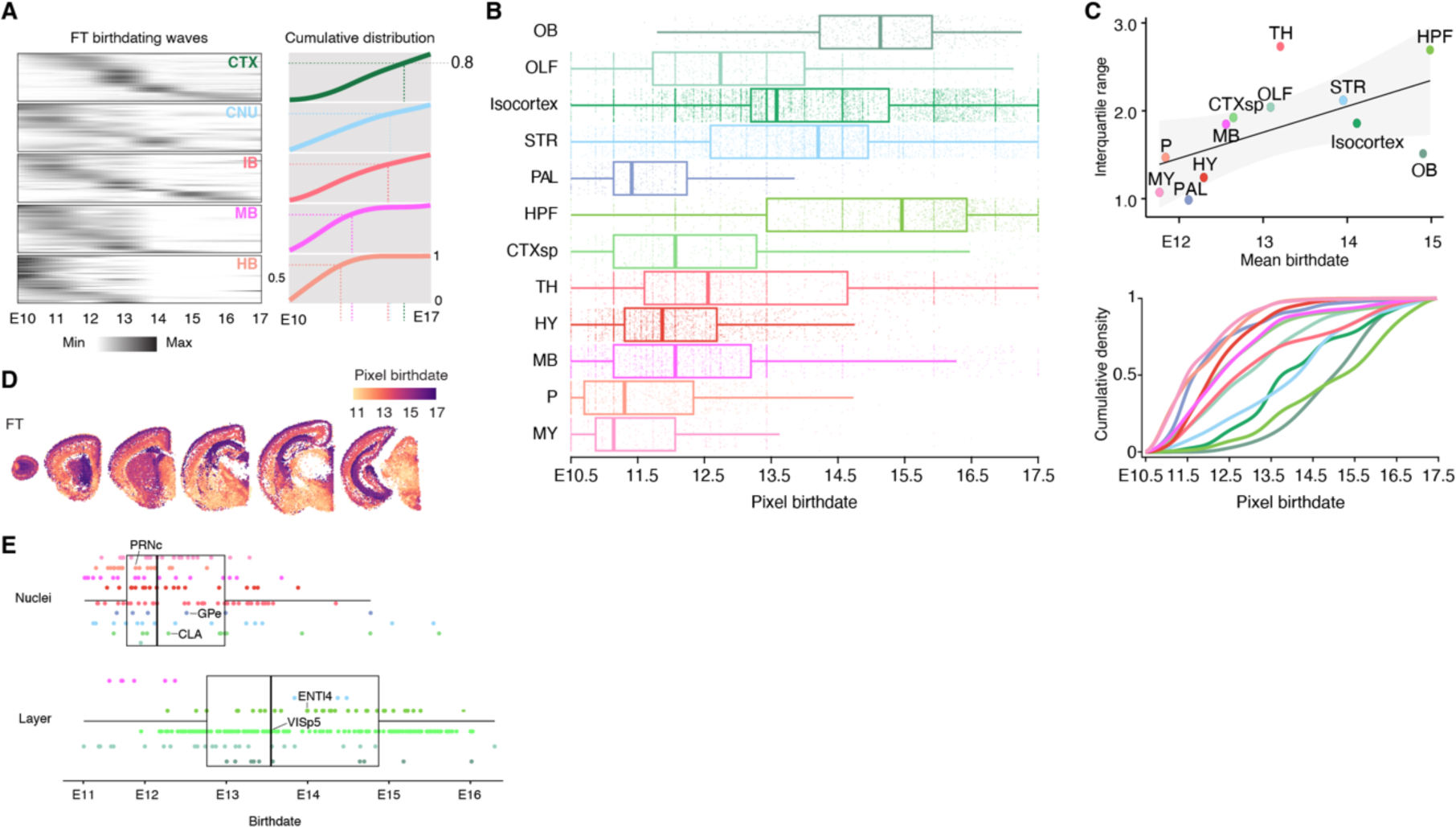
Brain-wide analysis of cellular birthdates with FT reveals rostral sustained-neurogenic and caudal transient-neurogenic regions. (A) FT birthdate-waves for all anatomical structures grouped by regions (left) and cumulative distribution of birthdate-waves (right). The time to reach 80% of born cells is highlighted by the dotted lines. (B) Superpixel birthdate split by supra-structures for the FT dataset. (C) FT mean birthdate against interquartile range of superpixel birthdate for supra-structures (top) and cumulative distribution of superpixel FT birthdate for supra-structures (bottom). (D) Superpixel birthdate maps for the 8 rostro-caudal sections of the FT dataset. (E) Birthdate of anatomical structures color-coded by their supra-structure and split into nuclear or laminar structure. OB: olfactory bulb; OLF: olfactory areas; STR: striatum; PAL: pallidum; HPF: Hippocampus; CTXsp: Cortical Subplate; TH: thalamus; HY: Hypothalamus; MB: Midbrain; P: Pons; MY: Medulla: CLA: claustrum; ENTl4: Entorhinal area layer 4; GPe: globus pallidus, external segment; PRNc: pontine reticular nucleus; VISp5: visual primary area layer 5.

**Figure S3.**
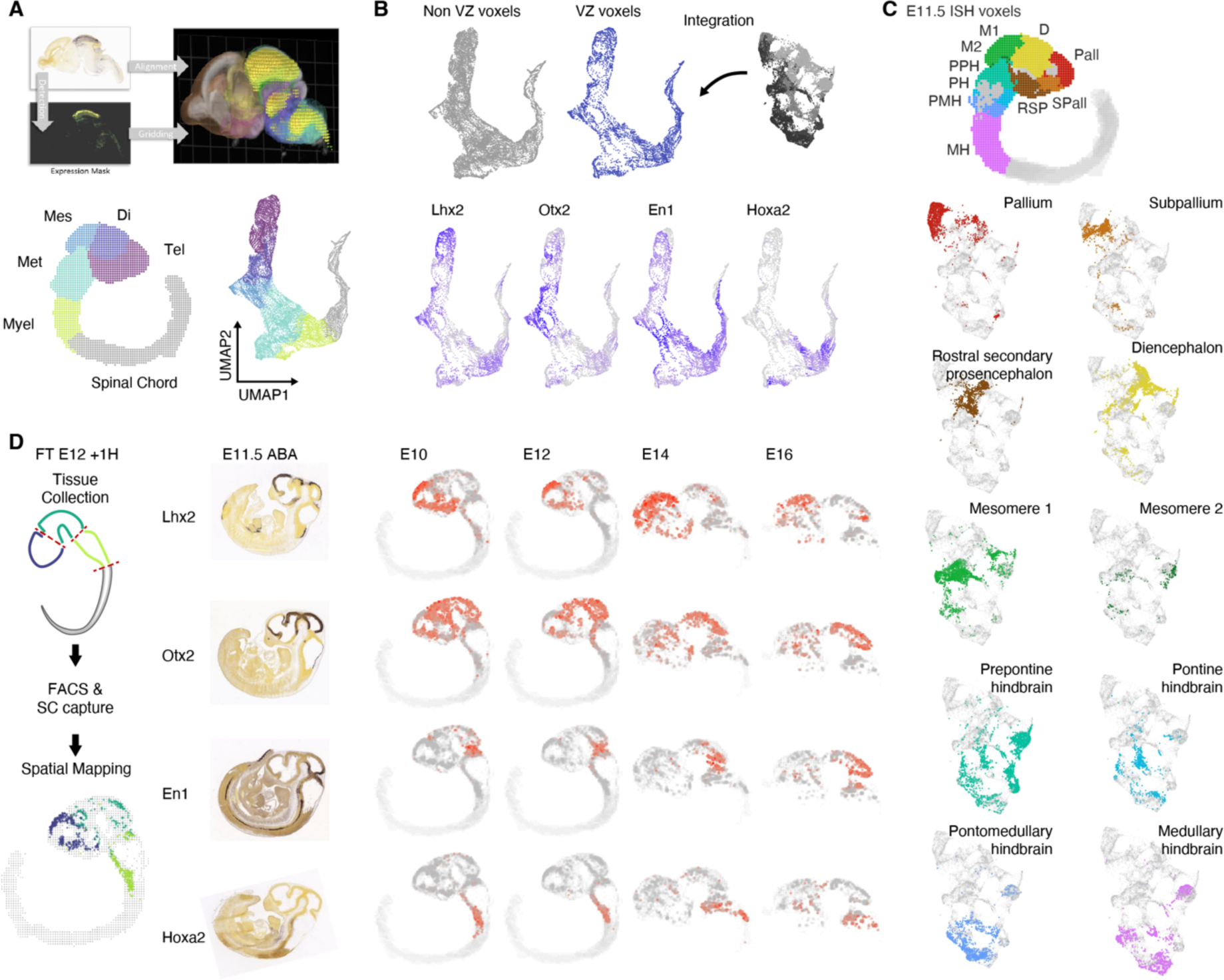
Spatial mapping of apical progenitors to voxel-based in situ hybridization gene expression data. (A) Schematic of the transformation of in situ hybridization (ISH) sections to 3D gene expression voxel dataset from the Allen Brain Atlas^17^ (top), Pseudo sagittal representation at E11.5 (bottom left) and UMAP presentation (bottom right) of ISH voxels color-coded by neural tube subdivisions. (B) UMAP plot of voxels outside of or within the VZ (top), and expression of known neural tube patterning genes in VZ voxels (bottom). (C) Voxels at E11.5 color-coded by neural tube subdivisions (top) and UMAP representation of cell mapping to these subdivisions (bottom). (D) Spatial mapping of cells collected from microdissected forebrain, midbrain and hindbrain segments at E12 (left). ISH sections of known neural tube patterning markers and expression in spatially map cells (right); image source: Allen Developmental Brain Atlas (developingmouse.brain-map.org). Tel: Telencephalon; Di Diencephalon; Met: Metencephalon; Myel: Myelencephalon; Pall: Pallium; Spall: Subpallium; RSP: rostral secondary prosencephalon; D: Diencephalon; M1: mesomere 1; M2: mesomere 2; PPH; Prepontine hindbrain; PH: Pontine hindbrain; PMH: Pontomedullary hindbrain; MH: Medullary hindbrain.

